# Changes over a 10-year Period in the Distribution Ranges and Genetic Hybridization of Three *Pelophylax* Pond Frogs in Central Japan

**DOI:** 10.1101/2025.05.28.656534

**Authors:** Shonosuke Shigeta, Tomoya Suzuki, Shohei Komaki, Koji Tojo

**Author notes:** These two authors contributed equally to this work (Shonosuke Shigeta, Tomoya Suzuki). **Corresponding author:** Koji Tojo, Department of Biology, Faculty of Science, Shinshu 21 University, Asahi 3-1-1, Matsumoto, Nagano 390-8621, Japan **E-mail address:**.

## Abstract

In this study, we focused on interspecial interactions in Central Japan where three closely related pond frog species, *Pelophylax nigromaculatus*, *Pelophylax porosus porosus* and *Pelophylax porosus brevipodus* (two species and two subspecies) are distributed and are in contact and/or overlap. The presence or absence of hybridization between these species was evaluated by comparing the sequences of the mitochondrial DNA Cyt-b (586-bp) and the nuclear gene Tyrosinase (747-bp) regions of each specimen collected. By comparing interspecific hybridization in the same area in both 2010 (119 specimens) and 2020 (152 specimens), we clarified population distributions and corresponding genetic dynamics over a 10-year period; there was no major change in the degree of interspecific hybridization between *P. nigromaculatus* and *P. p. brevipodus* in the Ina Basin, but it was revealed that the proportion of “pure”-bred *Pelophylax p. brevipodus* remained low. On the other hand, in the Matsumoto Basin, interspecific hybridization between *P. nigromaculatus* and *P. p. porosus* was evaluated to have progressed slightly. The proportion of “pure”-bred *Pelophylax p. porosus* declined to less than half, while the proportion of hybrid individuals showed a tendency to increase. Moreover, the dispersal of *P. nigromaculatus* was confirmed in areas along the Sai River, where only the pure-bred *P. p. porosus* was previously known to inhabit. Since the three frogs targeted in this study are all “Red List” species that are endangered, continuous monitoring of population structures and genetic dynamics is required.

## 1. Introduction

In Japan, a great number of floodplain environments have been lost due to river improvement projects (Tomita et al., 2020; Suzuki et al., 2023). Consequently, paddy fields spread across Japan have become habitats for many wetland aquatic organisms as an alternative to the lost floodplain habitats (Yamazaki et al., 2001; Kameyama et al., 2006). However, with the modernization of agriculture, the habitats of aquatic organisms have deteriorated due to the effects of pesticides and agricultural land development projects, and the number of paddy fields has decreased due to the conversion of agricultural fields from paddy fields to fields for uses other than rice (Katano et al., 2003; Yamamoto and Senga, 2012). Under such circumstances, the survival of many wetland aquatic species is in danger and they have been targeted for conservation as Red List species in Japan (Ministry of the Environment, Government of Japan, 2020). The *Pelophylax* pond frogs investigated in this study make up a typical species in crisis. *Pelophylax* pond frogs are one of the most common frog groups in Japan (Matsui and Maeda, 2018). They are widespread in river floodplains and are the most commonly seen frogs. Two species, *Pelophylax porosus* (Cope, 1868) and *Pelophylax nigromaculatus* (Hallowell, 1861), inhabit Japan. Of these, *P. porosus* is an endemic species to Japan, with the subspecies *P. porosus porosus* inhabiting Eastern Japan and the subspecies *P. porosus brevipodus* Western Japan (Fig. 1). The distribution of these subspecies is considered to be closely related to the formation history of the Japanese Archipelago (Komaki et al., 2015), and the distribution boundary of these subspecies is approximately along the Itoigawa-Shizuoka Tectonic Line, which is a large active fault (Sagiya et al., 2002). This geological tectonic line constitutes the western edge of the “Fossa Magna” region, which separated Northeast Japan from Southwest Japan by a deep strait 15–5 Ma (Fig. 1). On the other hand, *P. nigromaculatus* inhabits a wide area of the Japanese Archipelago, and also the continental region of East Asia (Fig. 1). Phylogenetic evolution and biogeographical studies of these *Pelophylax* pond frogs have already been conducted in considerable detail (Komaki et al., 2015). These studies suggest that *P. nigromaculatus* arrived in the Japanese Archipelago after *P. porosus* and expanded its distribution area. In addition, it has become clear that interspecific hybridization is occurring in the Japanese Archipelago (Nishioka et al., 1992; Komaki et al., 2012; Naito, 2012). Such increasing interspecific hybridization, along with the loss of habitats, are major concerns for the conservation of these species. Since hybrid males of *P. nigromaculatus* and *P. porosus* exhibit inferior reproductive capability (Moriya, 1960), extended reproductive interference induced by artificial disturbance of their habitat and subsequent population decline are also conservation concerns.

**Figure 1.**
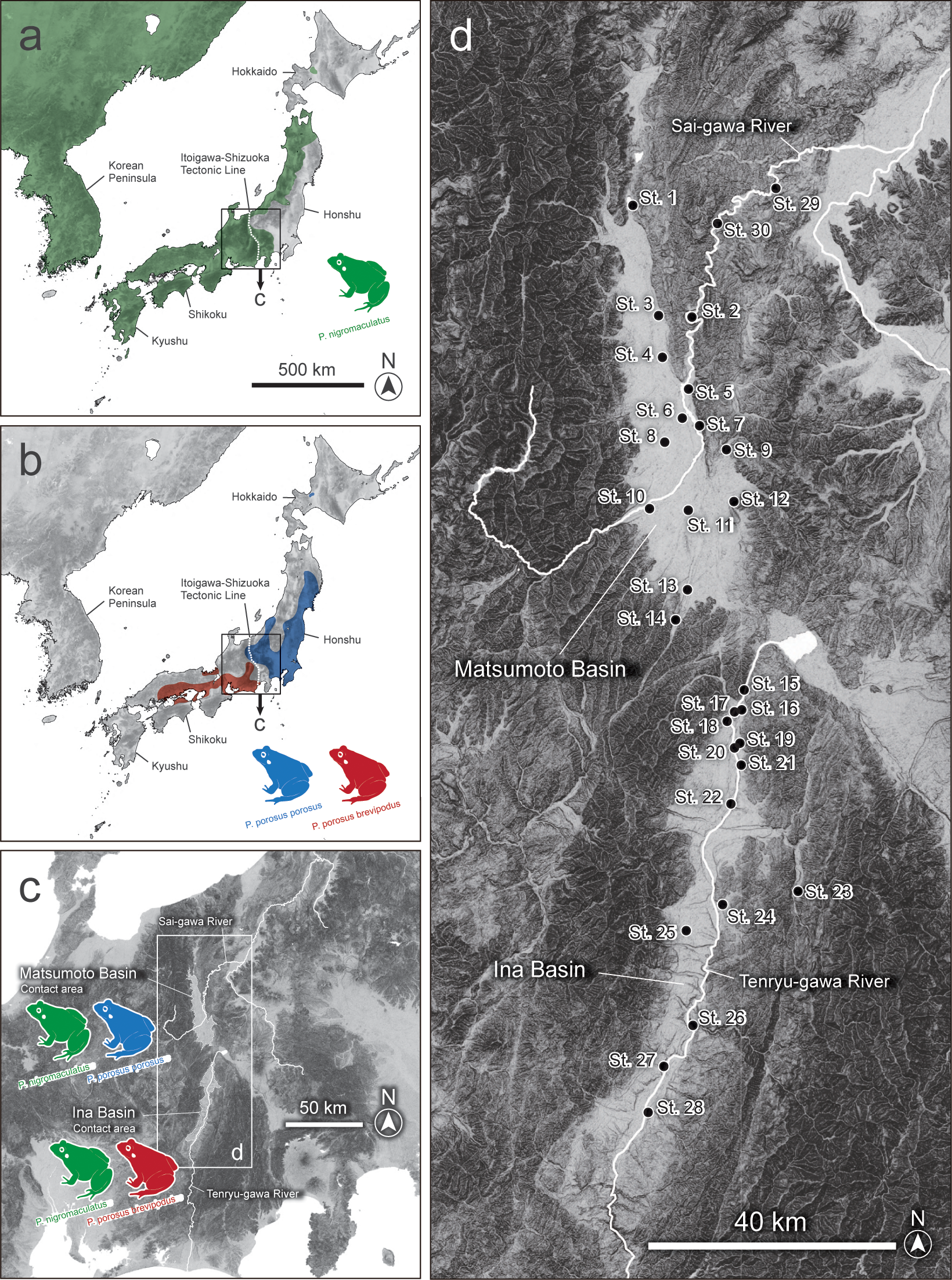
Survey sites in the Matsumoto Basin and the Ina Basin (Nagano Prefecture, Central Japan) of three species of *Pelophylax* pond frogs, and the distribution area to be analyzed in this study (Matsui and Maeda, 2018). a) Distribution range of *Pelophylax nigromaculatus*, b) distribution range of the species *Pelophylax porosus*, c) distribution range of the *Pelophylax* species, d) occurrences of species in the Matsumoto and Ina Basins, and e) a further enlarged map of the survey area (30 survey sites).

The study by Komaki et al. (2012) focused on the Matsumoto and Ina basins in Nagano Prefecture, which is a contact area between the two species (Fig. 1); the study produced detailed mesh-maps of these areas, and the habitat situation of the *Pelophylax* pond frogs within each mesh-map was investigated. In the study, in addition to the evaluation of morphological traits, a mitochondrial gene (mtDNA Cyt-b region) and nuclear gene (allozyme) analyses were performed to evaluate whether specimens were pure-bred *P. porosus*, *P. nigromaculatus*, or hybrid strains. In addition, genetic analyses of more than 100 frogs in each of the Matsumoto and Ina basins revealed that about 40% of the *Pelophylax* pond frogs in the Matsumoto basin and over 50% in the Ina basin were hybrids (Komaki et al., 2012). Comparing these results with the distribution of *P. porosus*, *P. nigromaculatus*, and the hybrid strain based on morphological characteristics in the 1980s, it became clear that the distribution area of *P. nigromaculatus* had expanded and also that the area in which the hybrid strain was found had expanded over about a 30-year period (Komaki et al., 2012).

Komaki et al. (2012) summarized the results of a survey conducted in 2010. In 2020, 10 years after the original survey, we conducted a follow-up survey and collected *Pelophylax* pond frogs using the same method as in the survey of Komaki et al. (2012). A mesh-map division of the same survey sites was conducted in the Matsumoto and Ina basins, a field survey was conducted, *Pelophylax* pond frogs were collected, and genetic analyses were conducted. By analyzing the mitochondrial and nuclear genes of 150 *Pelophylax* pond frog specimens collected from the Matsumoto basin and 130 specimens collected from the Ina basin, we differentiated between pure-bred *P. porosus* or *P. nigromaculatus* and a hybrid strain, as well as the change in their ratio in the 10 years from 2010 to 2020.

## 2. Materials and Methods

### 2.1 Sampling in the Matsumoto and Ina Basins

Based fundamentally on the sampling location information of Komaki et al. (2012), survey sampling of *Pelophylax* pond frogs in the Matsumoto and Ina basins was carried out (Fig. 1). The center of the previous survey sites were identified using GPS information, and sampling was carried out for a regular sampling duration (continuous effort for 30 min per study site) in the rice paddy fields and surrounding waterways centred on each location. Komaki et al. (2012) suggested that the distribution of *Pelophylax nigromaculatus* in the downstream direction of the Sai-gawa River had expanded, so two additional survey sites (sites No. 29 and 30) were newly set north of the Matsumoto basin. In addition, two new survey points (sites No. 26 and 28) were set to further refine the understanding of the distribution of *Pelophylax* pond frogs in the southern region of the Ina Basin. From May 15th to September 15th, 2020, 293 adult *Pelophylax* pond frogs were collected at all 30 sites shown in Figure 1.

All the collected *Pelophylax* pond frog specimens were identified by morphological characteristics based on Matsui and Maeda (2018) and Komaki and Tojo (2010), and photographs were taken from the dorsal and lateral views. Of these specimens collected for use in genetic analyses (293 specimens in total: Matsumoto Basin: 152 individuals, Ina Basin: 128 individuals, Sai-gawa downstream area: 13 individuals). Small tissue specimens were excised from the fourth toe of the hind limbs and stored in 100% EtOH. The GPS information, and elevation are shown in Table 1, and the number of samples collected and genetically analyzed is shown in Table 2.

**Table 1.** Study site information (Locality name, Latitude, Longitude, and Altitude).

| St. | Locality | Latitude (N) | Longitude (E) | Altitude (m) |
| --- | --- | --- | --- | --- |
| Matsumoto Basin |  |  |  |  |
| 1 | Taira, Omachi | 36.554536 | 137.839881 | 763 |
| 2 | Ikusaka | 36.426217 | 137.924294 | 497 |
| 3 | Ikeda, Ikeda | 36.428833 | 137.879408 | 619 |
| 4 | Aisome, Ikeda | 36.382028 | 137.882083 | 557 |
| 5 | Nakagawate, Akashina | 36.337542 | 137.924122 | 532 |
| 6 | Minami-hotaka, Toyoshina, Azumino | 36.314889 | 137.919222 | 536 |
| 7 | Tazawa, Toyoshina, Azumino | 36.300917 | 137.939611 | 550 |
| 8 | Misato, Azumino | 36.27475 | 137.889083 | 589 |
| 9 | Okada, Matsumoto | 36.277972 | 137.977056 | 710 |
| 10 | Hata, Matsumoto | 36.210333 | 137.867361 | 651 |
| 11 | Wada, Matsumoto | 36.207694 | 137.922417 | 612 |
| 12 | Kanda, Matsumoto | 36.213139 | 137.984778 | 604 |
| 13 | Seba, Shiojiri | 36.112333 | 137.922028 | 704 |
| 14 | Makino, Shiojiri | 36.082083 | 137.90975 | 759 |
| Ina Basin |  |  |  |  |
| 15 | Tatsuno, Tatsuno | 35.998278 | 138.004333 | 735 |
| 16 | Hiraide, Tatsuno | 35.975911 | 137.998853 | 711 |
| 17 | Miyaki, Tatsuno | 35.973306 | 137.990694 | 724 |
| 18 | Shinmachi, Tatsuno | 35.967028 | 137.980556 | 729 |
| 19 | Higashi-minowa, Minowa 1 | 35.936461 | 137.996569 | 708 |
| 20 | Higashi-minowa, Minowa 2 | 35.927589 | 137.988472 | 686 |
| 21 | Mikkamachi, Minowa | 35.915475 | 137.998258 | 676 |
| 22 | Minami-minowa | 35.865722 | 137.980917 | 657 |
| 23 | Nakao, Hase, Ina | 35.766944 | 138.079694 | 848 |
| 24 | Higashi-ina, Komagane | 35.750639 | 137.975 | 617 |
| 25 | Akaho, Komagane | 35.719861 | 137.913778 | 723 |
| 26 | Katsurashima, Nakagawa | 35.606583 | 137.929833 | 470 |
| 27 | Yoshida, Takamori | 35.562028 | 137.891361 | 426 |
| 28 | Takagi | 35.508806 | 137.864583 | 403 |
| The downstream basin of the Sai-gawa River |  |  |  |  |
| 29 | Shinshu-shinmachi, Nagano | 36.575167 | 138.052056 | 413 |
| 30 | O-okakou, Nagano | 36.536639 | 137.965139 | 465 |

**Table 2.** The number of individuals that were identified as *P. nigromaculatus*, *P. porosus porosus*, *P. porosus brevipoda*, and hybrid, based on morphological characteristics, mtDNA Cyt-b, and nDNA Tyrosinase sequences in each study site.

| St. | Locality | Sample size | Morphological characteristics |  |  |  | mtDNA Cyt-b |  |  | nDNA Tyrosinase |  |  |  |
| --- | --- | --- | --- | --- | --- | --- | --- | --- | --- | --- | --- | --- | --- |
|  |  |  | N | P | B | H | N | P | B | N | P | B | H |
| Matsumoto Basin |  |  |  |  |  |  |  |  |  |  |  |  |  |
| 1 | Taira, Omachi | 19 | 15 |  |  | 4 | 19 |  |  | 6 |  |  | 13 |
| 2 | Ikusaka | 18 | 17 |  |  | 1 | 12 | 6 |  | 8 |  |  | 10 |
| 3 | Ikeda, Ikeda | 12 |  | 12 |  |  |  | 12 |  |  | 4 |  | 8 |
| 4 | Aisome, Ikeda | 11 |  | 10 |  | 1 |  | 11 |  |  | 5 |  | 6 |
| 5 | Nakagawate, Akashina | 9 | 9 |  |  |  | 9 |  |  | 6 |  |  | 3 |
| 6 | Minami-hotaka, Toyoshina, Azumino | 9 | 9 |  |  |  | 8 | 1 |  | 8 |  |  | 1 |
| 7 | Tazawa, Toyoshina, Azumino | 9 | 9 |  |  |  | 7 | 2 |  | 7 |  |  | 2 |
| 8 | Misato, Azumino | 9 | 2 | 8 |  |  | 3 | 6 |  | 3 |  |  | 6 |
| 9 | Okada, Matsumoto | 12 |  | 12 |  |  |  | 12 |  | 7 |  |  | 5 |
| 10 | Hata, Matsumoto | 4 | 4 |  |  |  | 3 | 1 |  | 3 |  |  | 1 |
| 11 | Wada, Matsumoto | 8 | 8 |  |  |  | 8 |  |  | 8 |  |  |  |
| 12 | Kanda, Matsumoto | 12 | 8 | 4 |  |  | 9 | 3 |  | 5 |  |  | 7 |
| 13 | Seba, Shiojiri | 10 | 9 |  |  | 1 | 9 | 1 |  | 4 |  |  | 6 |
| 14 | Makino, Shiojiri | 10 | 10 |  |  |  | 3 | 7 |  | 5 |  |  | 5 |
| Ina Basin |  |  |  |  |  |  |  |  |  |  |  |  |  |
| 15 | Tatsuno, Tatsuno | 8 | 8 |  |  |  | 8 |  |  | 8 |  |  |  |
| 16 | Hiraide, Tatsuno | 12 | 11 |  | 1 |  |  |  |  | 7 |  | 4 | 1 |
| 17 | Miyaki, Tatsuno | 12 | 5 |  | 6 | 1 | 8 |  | 4 | 4 |  | 3 | 5 |
| 18 | Shinmachi, Tatsuno | 3 | 3 |  |  |  | 3 |  |  | 3 |  |  |  |
| 19 | Higashi-minowa, Minowa 1 | 11 | 4 |  | 2 | 5 | 6 |  | 5 | 3 |  | 3 | 5 |
| 20 | Higashi-minowa, Minowa 2 | 7 | 1 |  | 7 |  | 3 |  | 5 | 1 |  | 4 | 3 |
| 21 | Mikkamachi, Minowa | 14 | 5 |  | 8 | 1 | 7 |  | 7 | 3 |  | 6 | 5 |
| 22 | Minami-minowa | 10 | 1 |  | 9 |  | 4 |  | 6 | 1 |  | 8 | 1 |
| 23 | Nakao, Hase, Ina | 3 | 3 |  |  |  | 3 |  |  | 3 |  |  |  |
| 24 | Higashi-ina, Komagane | 10 | 10 |  |  |  | 10 |  |  | 5 |  |  | 5 |
| 25 | Akaho, Komagane | 11 | 11 |  |  |  | 10 |  | 1 | 11 |  |  |  |
| 26 | Katsurashima, Nakagawa | 10 | 10 |  |  |  | 10 |  |  | 8 |  |  | 2 |
| 27 | Yoshida, Takamori | 9 | 1 |  | 8 |  | 3 |  | 6 | 1 |  | 5 | 3 |
| 28 | Takagi | 8 | 8 |  |  |  | 5 |  | 3 | 7 |  |  | 1 |
| The downstream basin of the Sai-gawa River |  |  |  |  |  |  |  |  |  |  |  |  |  |
| 29 | Shinshu-shinmachi, Nagano | 10 | 6 | 3 |  | 1 | 10 |  |  | 4 |  |  | 6 |
| 30 | O-okakou, Nagano | 3 | 2 | 1 |  |  | 3 |  |  | 2 |  |  | 1 |
|  | Total | 293 | 189 | 50 | 41 | 15 | 183 | 62 | 37 | 141 | 9 | 33 | 111 |

N: *Pelophylax nigromaculatus*, P: *Pelophylax porosus porosus*, B: *Pelophylax porosus brevipoda*, H: hybrid.

### 2.2 Genetic analyses

We extracted total genomic DNA from ethanol-preserved tissue specimens and purified it with a DNeasy Blood and Tissue Kit (Qiagen, Hilden, Germany), according to the manufacturer’s instructions.

We used total DNA to amplify DNA fragments of mitochondrial DNA (mtDNA), cytochrome b (Cyt-b) (610-bp) by polymerase chain reaction (PCR) with these sets of primers: L14850 (5′-TCT CAT CCT GAT GAAACT TTG GCT C-3′) and H15410 (5′-GTC TTT GTA GGA GAA GTA TGG-3′) (Tanaka et al., 1996). The PCR protocol was: 94°C for 1 min, 35 cycles (94°C for 1 min, 49°C for 1 min, 72°C for 1 min), 72°C for 7 min. We purified PCR products with Illustra ExoProStar (GE Healthcare, Buckinghamshire, UK).

We used total genomic DNA to amplify DNA fragments of the nuclear DNA (nuDNA) Tyrosinase (747-bp) by PCR with these sets of primers: TyrF_Rana (5’-TTC CCC TTG GTT TGT TTG AG-3’) (Iwasawa et al., in preparation) and TyrIG (5’-TGC TGG GCR TCT CTC CAR TCC CA-3’) (Bossuyt and Milinkovitch 2000). The PCR proximal protocol was 94°C for 1 min, 35 cycles (94°C for 1 min, 49°C for 1 min, 72°C for 1 min) and 72°C for 7 min. We purified PCR products with Illustra ExoProStar (GE Healthcare, Buckinghamshire, UK).

We sequenced purified DNA fragments directly with an automated method using a BigDye Terminator Cycle Sequencing Kit v. 1.1 (Applied Biosystems, Foster City, CA, USA) on an automated DNA sequencer (ABI 3130xl DNA Analyzer; Applied Biosystems). Sequence alignment and editing were performed for each gene separately with MEGA v. 7 (Kumar et al., 2016) and CLC Workbench software (CLC bio, Aarhus, Denmark). We have submitted all of the sequence data to GenBank (accession numbers: ######–########).

The haplotype of the Cyt-b region was determined using DNasP v6, (Rozas et al., 2017) and haplotype networks were constructed using the median-joining method in PopART (Leigh & Bryant, 2015). We used a data set combining sequence data from 2010 and 2020 to determine the haplotypes likely to occur in the study area and then compared the actual haplotypes that occurred for each year. The sequence data used by Komaki et al. (2012) was used to determine haplotypes, and we created haplotype networks for the 2010 and 2020 combined data set and also for each of the distinct 2010 (GenBank accession numbers: AB686619–41) and 2020 data sets (######–##).

As for the sequencing results of the Tyrosinase region, we also tested for the presence or absence of double peaks in the waveform data, particularly at species(subspecies)-specific variant sites (Table 3). Variable nucleotide sites were determined by aligning sequences without double peaks and identifying positions where variations were observed between species or subspecies, while all sequences within a species shared the same nucleotide. Of the 293 individuals analyzed, no double peaks were detected in 119 specimens, and so they were evaluated as pure-bred specimens of either species, consistent with evaluations based on morphology and mitochondrial DNA.

**Table 3.** Variable nucleotide sites identified in the 747 bp Tyrosinase sequence and used for species identification. Nucleotides highlighted in gray indicate positions that differ from those in other species (or subspecies). Numbers refer to the position beginning from the 5’ end.

| Species, subspecies | Site number |  |  |  |  |  |  |  |  |  |
| --- | --- | --- | --- | --- | --- | --- | --- | --- | --- | --- |
|  | 13 | 32 | 46 | 74 | 81 | 219 | 397 | 532 | 636 | 708 |
| *Pelophylax nigromaculatus* | A | T | A | C | C | C | A | G | T | A |
| *Pelophylax porosus porosus* | A | C | G | T | T | G | T | G | C | G |
| *Pelophylax porosus brevipodus* | G | C | G | T | T | G | A | A | C | G |

Regarding nuclear DNA analysis, a study by Komaki et al. (2012) in which they assessed the hybridization status of the two species in the 2010 survey, used allozyme analysis to determine whether the individuals were of the pure or hybrid breeding lineages of both species. In the 2020 survey, i.e., 10 years after that, hybridization status was assessed by sequencing the nuclear DNA tyrosinase region rather than by allozyme analysis. Based on our preliminary analysis of the nuclear DNA tyrosinase region using specimens from areas in which only one species inhabit, away from the distribution boundary between *P. porosus* and *P. nigromaculatus*, it was revealed that the two species can be easily distinguished from each other based on their genetic differentiation. Therefore, by analyzing the nuclear DNA tyrosinase region, specimens evaluated as “hybrid” in these data sets can be considered to be of hybrid individuals with high confidence, as hybrid strains will possess sequences from both species in the heterozygous state, or there will be discrepancies between mitochondrial DNA and nuclear DNA. If backcrossing has continued for a long period of time after past hybridization, it may no longer be possible to evaluate the lineage as being hybrid. However, this limitation applies to both allozyme and tyrosinase based analyses. In particular, since the focus of this study is on the increasing trend toward hybrid individuals, such problems of identifying hybrid individuals with certainty, even if there is a possibility that the proportion of hybrid individuals may be underestimated, there is no way that it could be overestimated. Therefore, the differences in using the differing nuclear DNA analysis methods for specimens collected in 2010 and 2020 are not considered to be problematic for the purposes of this study.

### 2.3 Calculation of the proportion of genetically pure-bred individuals (non-hybrid individuals) in each basin

In all of the morphological trait and inheritance analysis results (mitochondrial DNA and nuclear DNA analysis results), the proportion of individuals that clearly showed pure-bred lineage (non-hybrid individuals) was calculated. Thereafter, we evaluated the statistical significance of the difference in the proportion of pure-bred individuals vs hybrid individuals estimated between 2010 and 2020 (i.e., a chi-square test was performed on each species). In the analysis of the Matsumoto Basin, the proportions were compared exclusively within the basin, excluding specimens from the Sai-gawa downstream area (sites No. 29 and No. 30), as in Komaki et al. (2012). In the analysis of the Ina Basin, the two cases for which the data of the two newly added research sites in the 2020 survey were included, and the case where the data was not included, were evaluated separately. Bonferroni correction for multiple testing was used for results with significant *p* values. Corrected values (*p*) < 0.05 were considered significant.

### 2.4 Environmental analyses for the two basins over the last few decades

First, in the Matsumoto and Ina Basins, which are the target of this study, observational data from the Matsumoto Basin (Matsumoto Meteorological Observatory) and the Ina Basin (Iida Meteorological Observatory) for the 50 years from 1970 to 2020 was obtained based on public data (publicly available data from the Japan Meteorological Agency, https://www.jma.go.jp/jma/menu/menureport.html). Then, the temperature data of these 50 years (the trend in monthly average daily maximum temperatures of over a 50-year period; the trend in monthly average daily minimum temperatures; the trend in monthly average daily temperatures) and precipitation data (trends in monthly precipitation over the 50-year period) were analyzed. “Seasonal and Trend decomposition using, and “Remainder” of temperatire of decomposition using the software LOESS (Cleveland et al., 1990) was performed. The long-term trends of value fluctuations (trend components) obtained using the “stl” function in the R software were analyzed.

We also investigated the land use situation in the areas covered by the survey sites in the Matsumoto and Ina Basins. Regarding the analysis of land usase, we used GIS data published by the Ministry of Land, Infrastructure, Transport and Tourism (https://nlftp.mlit.go.jp/ksj/index.html), and used the data from the survey year closest to our current survey years (2010 and 2020). In addition, our previous study (Komaki et al., 2012) had referred to the 1878–1985 distribution data of the *Pelophylax* pond frog species group by Shimoyama (1986) and discussed the progress of hybridization of the *Pelophylax nigromaculatus* frog species group over the 30 years from 1985 to 2010, so we also analyzed the land use data for 1997. Excluding forests, the areas of the next three largest exclusive land use category in both basins were calculated for each survey year.

## 3. Results

### 3.1 Results of field sampling in 2020, and results of morphological characteristics and genetic analyses

In our 2020 survey, a total of 280 *Pelophylax* pond frog specimens were collected, from the Matsumoto Basin (152 samples) and the Ina Basin (128 samples) (Table 2, Figs 2, 3). In addition, 13 *Pelophylax* pond frog specimens were collected at the two survey sites along the Sai-gawa River newly added in the 2020 survey (Table 2, Fig. 2). Figure 2 shows the results of the morphological characteristics and the genetic analyses of all of the individuals analyzed.

**Figure 2.**
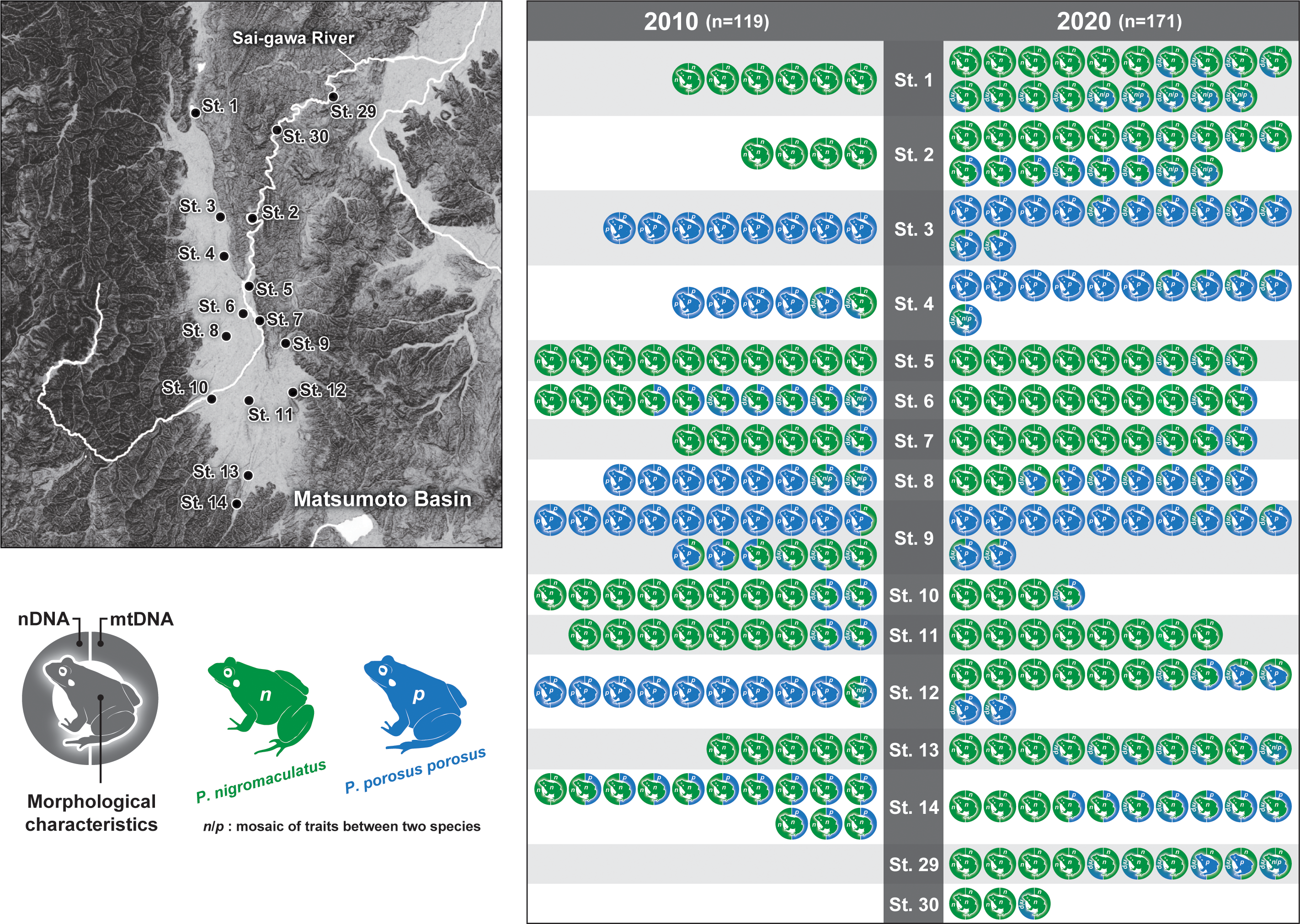
Changes over 10 years in the population and genetic structures (*i.e.*, nuclear DNA and mitochondrial DNA) of two *Pelophylax* pond frogs in the Matsumoto Basin. Evaluation results of morphological characteristics and the results of our genetic analyses at 14 sites surveyed in both 2010 and 2020 and two sites newly added in 2020. The color scheme of the frog marked in the center of the legend indicates the result of species identification by morphological characteristics. The color scheme of the right, semicircular marks, represents species identification based on mitochondrial DNA, while that of the left represents identification based on nuclear DNA. For each color scheme, the gradations in color indicate a mosaic of traits from the parent species.

**Figure 3.**
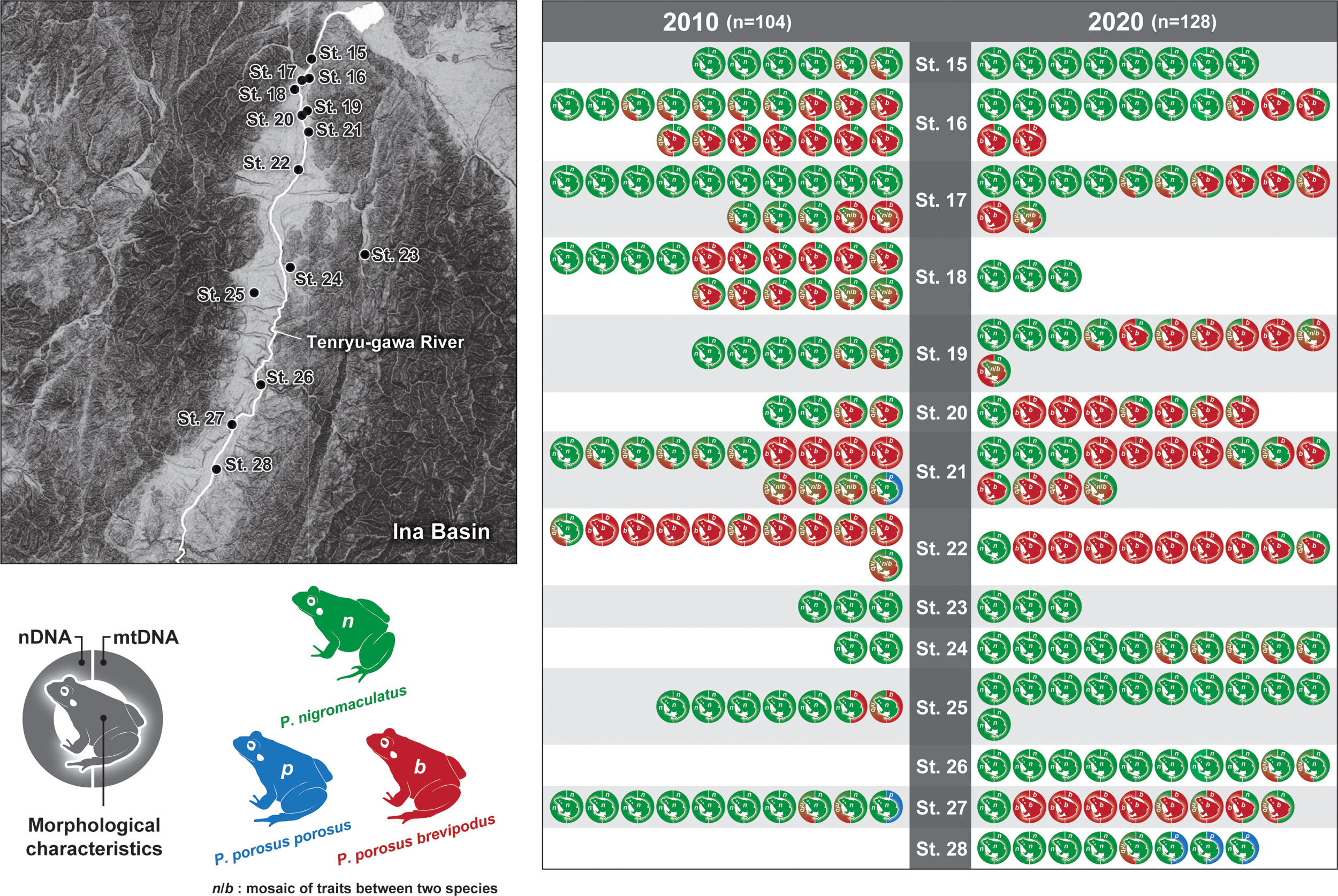
Changes over 10 years in the population and genetic structures (*i.e.*, the nuclear DNA and the mitochondrial DNA) of two *Pelophylax* pond frogs in the Ina Basin. Evaluation results of morphological characteristics and the results of our genetic analyses at 14 sites surveyed in both 2010 and 2020. The explanation for the color schemes is the same as in the legend of Figure 2.

Regarding the presentation of these results, the color scheme of the frog marked in the center of the legend indicates the result of species identification by morphological characteristics (Figs 2, 3). Specimens with a two-tone colored frog indicate individuals whose morphological characteristics suggested the possibility of crossbreeding of two *Pelophylax* pond frog species (Figs 2, 3). In addition, there are two semicircular marks forming a ring on the left and right of each of the central frog marks (Figs 2, 3). The color scheme of the semicircle mark on the right side of these indicates the result of species identification based on the mitochondrial DNA. Since mitochondrial DNA is maternally inherited, sequence data for the Cyt-b region is always retained within the lineage of each species. On the other hand, the color scheme of semicircular marks on the left indicates the results of species identification based on nuclear DNA. In terms of nuclear DNA sequences, it was suggested that both pure-bred strains each constitute distinct phylogenetic clades. On the other hand, since the *Pelophylax* pond frogs of hybrid individuals have sequences of both species in a mosaic manner, these hybrid individuals are shown with semi-circle marks in two-tone color. That is, if all the marks of each analyzed specimen have the same coloration, it indicates that they were all pure-bred specimens. On the other hand, specimens with a two-tone color mark are of a hybrid individuals.

The analysis yielded 22 haplotypes within the available Cyt-b region (493 bp) in the combined data set (Fig. 4a). The haplotypes were assigned to the 2010 and 2020 datasets based on the haplotypes that were identified in the combined dataset. 19 haplotypes were identified from the 2010 data set, and 11 haplotypes were identified from the 2020 data set (Figs 4b, 4c). Both the 2010 and 2020 data sets clearly distinguished each species (subspecies) into three haplogroups. In 2010, several minor haplotypes consisting of a small number of individuals were identified, but some of these minor haplotypes (n3, n5, n7, n8, n9, n10, n11, n12, n13, n15, p2, b2) were not detected in 2020, while some minor haplotypes (n16, n17, n18, n19) were newly identified (Figs 4b, 4c).

**Figure 4.**
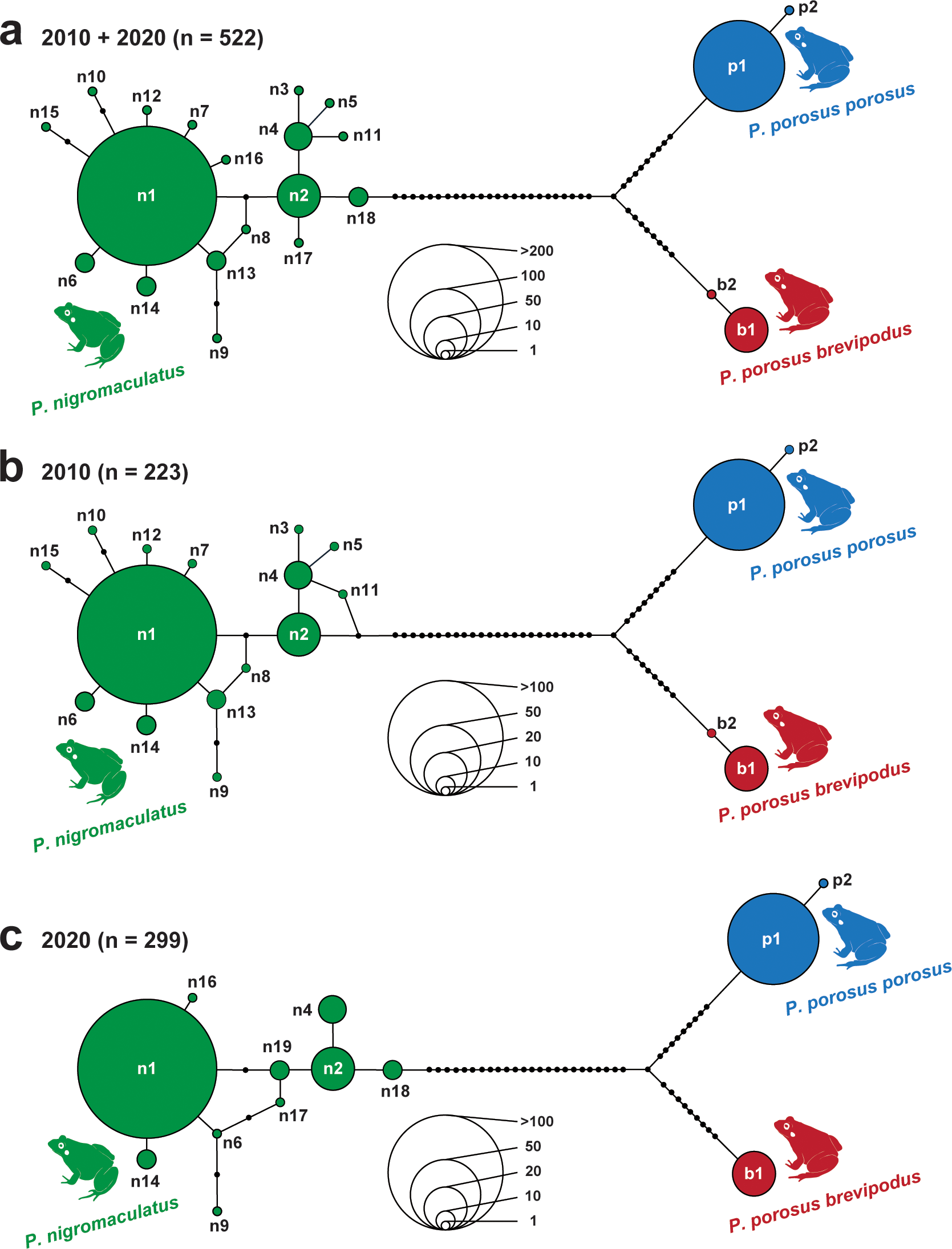
Median-joining haplotype network based on 493-bp of the mtDNA Cyt-b region obtained from surveys in 2010 and 2020. The size of the circles is proportional to the number of samples/sequences. The filled small circle(s) on the branch(es) connecting the circles indicate(s) the number of base substitutions between haplotypes. However, one filled small circle for a single substitution is omitted. (a) Network combining samples from both years (2010 and 2020 surveys). (b) Network generated from the 2010 survey data. (c) Network generated from the 2020 survey data. “n” is the number of samples used to create each network. The value of “n” indicates the number of samples used to construct each network.

### 3.2 Changes in the proportion of pure-bred lineages vs hybrid individuals of two *Pelophylax* pond frog species in the Matsumoto Basin over 10 years (2010–2020)

In the 10 years from 2010 to 2020, the proportion of pure-bred *Pelophylax nigromaculatus* decreased from 39.8 to 36.2%, and the proportion of pure-bred *Pelophylax porosus porosus* also decreased from 29.8% to 11.2% (Table 4, Figs 2, 5). On the other hand, the hybrid descendants of both species increased from 29.8% to 52.6% (Table 4, Figs 2, 5). At the four survey sites (No. 1, 2, 5 and 13), although only pure-bred *P. nigromaculatus* specimens were observed in 2010, hybrid descendants were also observed in 2020 (Table 4, Figs 2, 5). In addition, at site No. 3, where only pure-bred *P. p. porosus* specimens were observed in 2010, many hybrid individuals were also observed in 2020 (Table 4, Figs 2, 5). Among these hybrid individuals, the proportion of the type with the mitochondrial DNA of *P. nigromaculatus* increased from 22.2% to 43.8% (Table 4, Figs 2, 5).

**Table 4.** The number of individuals and its proportion of the “pure” *P. nigromaculatus*, *P. porosus porosus*, *P. porosus brevipoda*, and hybrid collected in the Matsumoto Basin and Ina Basin excluding the Sai-gawa downstream area in 2010 and 2020. “*” indicates a significant difference (*p*=0.05) between the proportion in 2010 and 2020.

|  | Number of "pure" individuals |  |  | Hybrid | Total |
| --- | --- | --- | --- | --- | --- |
|  | N | P | B |  |  |
| Matsumoto Basin |  |  |  |  |  |
| 2010 | 49 (41.1%) | 36 (30.3%) | 0 | 34 (28.6%) | 119 |
| 2020 | 55 (36.2%) | 17 (11.2%)* | 0 | 80 (52.6%)* | 152 |
| Ina Basin |  |  |  |  |  |
| 2010 | 44 (42.3%) | 0 | 9 (8.7%) | 51 (49.0%) | 104 |
| 2020 | 62 (48.4%) | 0 | 20 (15.6%) | 46 (35.9%) | 128 |

N: *Pelophylax nigromaculatus*, P: *Pelophylax porosus porosus*, B: *Pelophylax porosus brevipoda*.

**Figure 5.**
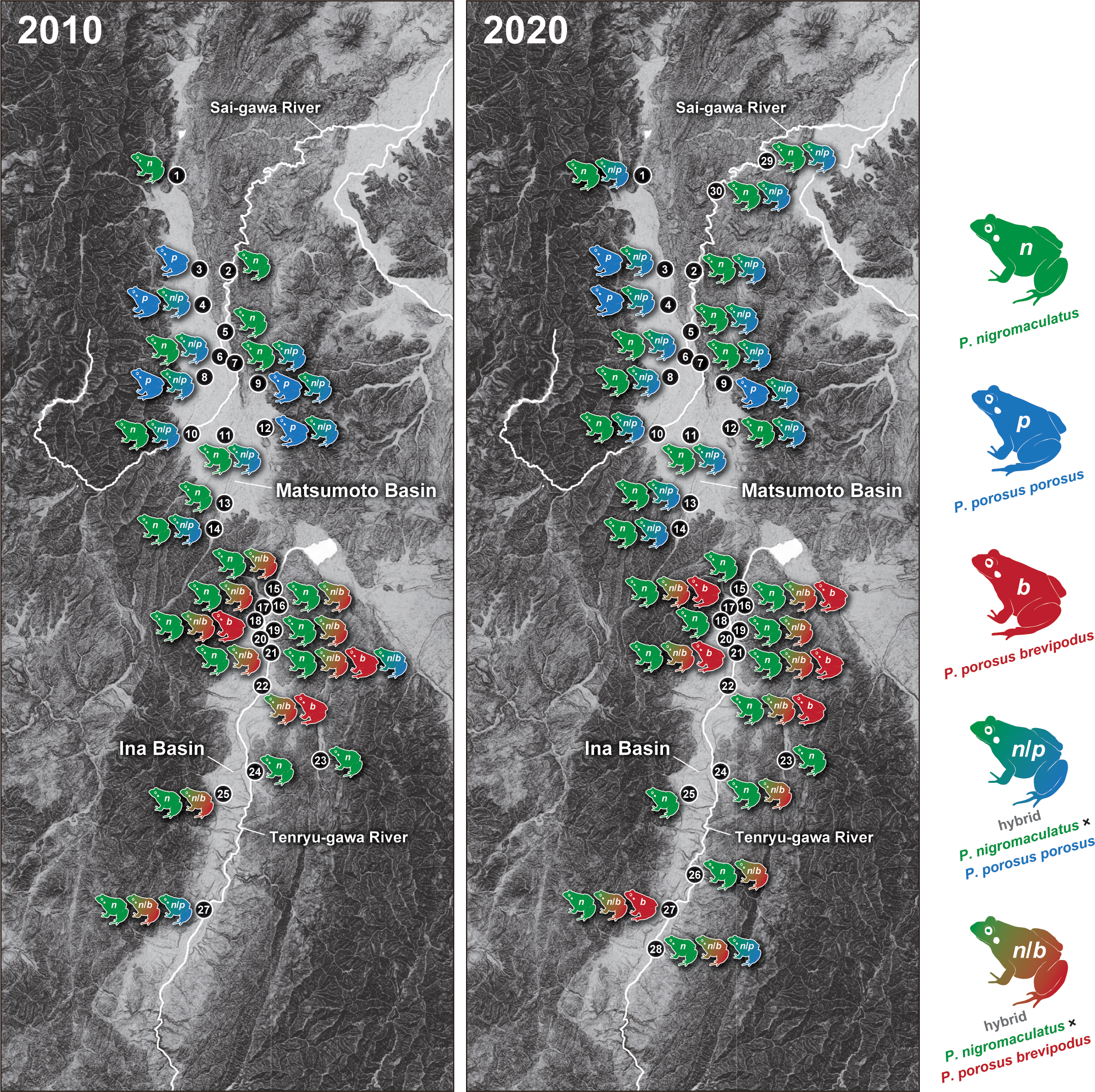
Distributions of pure-bred and hybrid strains of the three species/subspecies analyzed and their changes over the period 2010–2020 at each survey site in the Matsumoto Basin and the Ina Basin (Nagano Prefecture, Central Japan). This figure does not reflect the proportion of the pure-bred/hybrid individuals detected by genetic analyses. If even one individual of a specific type was detected, it was treated as “present”.

### 3.3 Changes in the proportion of pure-bred lineages vs hybrid individuals of two *Pelophylax* pond frog species in the Ina Basin over 10 years (2010–2020)

In the 10 years from 2010 to 2020, the proportion of pure-bred *Pelophylax nigromaculatus* increased from 39.6% to 45.5%, while the proportion of pure-bred *Pelophylax porosus brevipodus* also increased from 8.1% to 18.2% (Table 4, Figs 3, 5). On the other hand, the proportion of hybrid individuals decreased from 52.3% to 36.4% (Table 4, Figs 3, 5). When including the data of the two new sites, No. 26 and 28, added in the 2020 survey, the ratio of pure-bred *P. nigromaculatus* was 48.4%, the proportion of hybrid individuals was 15.6%, and the ratio of hybrid individuals was 35.9% (Table 4, Figs 3, 5). No survey sites were found in either of the 2010 or 2020 studies in which only pure-bred *P. p. brevipodus* were detected (Table 4, Figs 3, 5). At site No. 23, only pure-bred *P. nigromaculatus* specimens were detected in both 2010 and 2020 (Table 4, Figs 3, 5). In addition, at the No. 15 and 25 survey sites, both pure-bred *P. nigromaculatus* and hybrid individuals were detected in 2010, while only pure-bred *P. nigromaculatus* specimens were detected in 2020 (Table 4, Figs 3, 5).

In addition to our 10-year interval surveys in 2010 and 2020, we also analyzed available open data on climate change and land use in both basins from the first survey (1980s) conducted in a previous study (Shimoyama, 1986) to the present. Although no clear relationship was found with the changes in the distribution area of the two *Pelophylax* pond frog species or the progress of interspecific hybridization, the results of the analyses of environmental factors are shown in Figures S1 and S2; related land use statistics are provided in Table S1.

### 3.4 Analysis results for the Sai-gawa River Basin (downstream area of the Matsumoto Basin) newly added in the 2020 survey

The downstream basin of the Sai-gawa River (sites No. 29 and No. 30) was traditionally thought to be inhabited only by *Pelophylax porosus porosus* (Maeda and Matsui 2003), so it was not included in the previous survey area in 2010 (Komaki et al., 2012). However, when new survey sites (sites No. 29 and No. 30) were added in the 2020 survey, pure-bred *Pelophyalax porosus porosus* were not detected; even so, the presence of pure-bred *Pelophylax nigromaculatus* was confirmed (Fig. 2).

### 3.5 Observation of *Pelophylax porosus porosus* haplotypes in the Ina Basin

In our previous survey conducted in 2010 at survey sites No. 21 and No. 27 in the Ina Basin, which are not considered to be within the distribution area of *Pelophylax porosus porosus*, one specimen was collected from each site exhibiting the mtDNA haplotype of *Pelophylax porosus porosus* (not *Pelophylax porosus brevipodus*; of which the morphological characteristics are the same as those of *Pelophylax nigromaculatus*) (Figs 3, 5). In the current survey in 2020, a specimen showing similar traits was detected at an adjacent site, No.28 (i.e., morphologically *Pelophylax nigromaculatus*, but of the *Pelophylax porosus porosus* type according to its mtDNA) (Figs 3, 5).

### 3.6 Dynamics of *Pelophylax* pond frogs in two basins over a 10-year period

Of all specimens excluding the Sai-gawa downstream area analyzed in—2020, genetically pure-bred *Pelophylax nigromaculatus* accounted for 117 individuals, or 41.8%, whilst genetically pure-bred *Pelophylax porosus* accounted for 13.2%, with 37 individuals (Table 4). Meanwhile, the hybrid specimens accounted for 45.0%, with 126 individuals (Table 4).

There was no significant change in the proportion of pure-bred *Pelophylax nigromaculatus* over the decade from 2010 to 2020 in either the Matsumoto Basin or the Ina Basin (Table 4). However, it was shown that the percentage of hybrid individuals (*Pelophylax nigromaculatus* × *Pelophylax porosus porosus*) increased (*p*<0.05) in the Matsumoto Basin, while the proportion of *P. p. porosus* declined (*p*<0.05) (Table 4; Fig. 6). On the other hand, in the Ina Basin, the overall percentage of hybrid individuals (*Pelophylax nigromaculatus* × *Pelophylax porosus brevipodus*) decreased when newly added survey sites were included in the 2020 survey (Table 4; Fig. 6). It also showed an increased proportion of genetically pure-bred *Pelophylax porosus brevipodus*, even when not including the newly added sites. However, changes in the proportions of hybrid and purebred individuals in the Ina Basin were not statistically significant according to the chi-square test. In the 2010 survey, pure-bred *P. p. brevipodus* accounted for 8.7% (9/104), while in the 2020 survey at the same site it was 18.0% (20/111), and in the survey of all sites including the newly added sites (St. 26, 28), it was 15.6% (20/128). Therefore, there was no indication that the proportion of pure-bred *Pelophylax porosus brevipodus* had decreased during the last 10 years in the Ina Basin (Table 4; Fig. 6).

**Figure 6.**
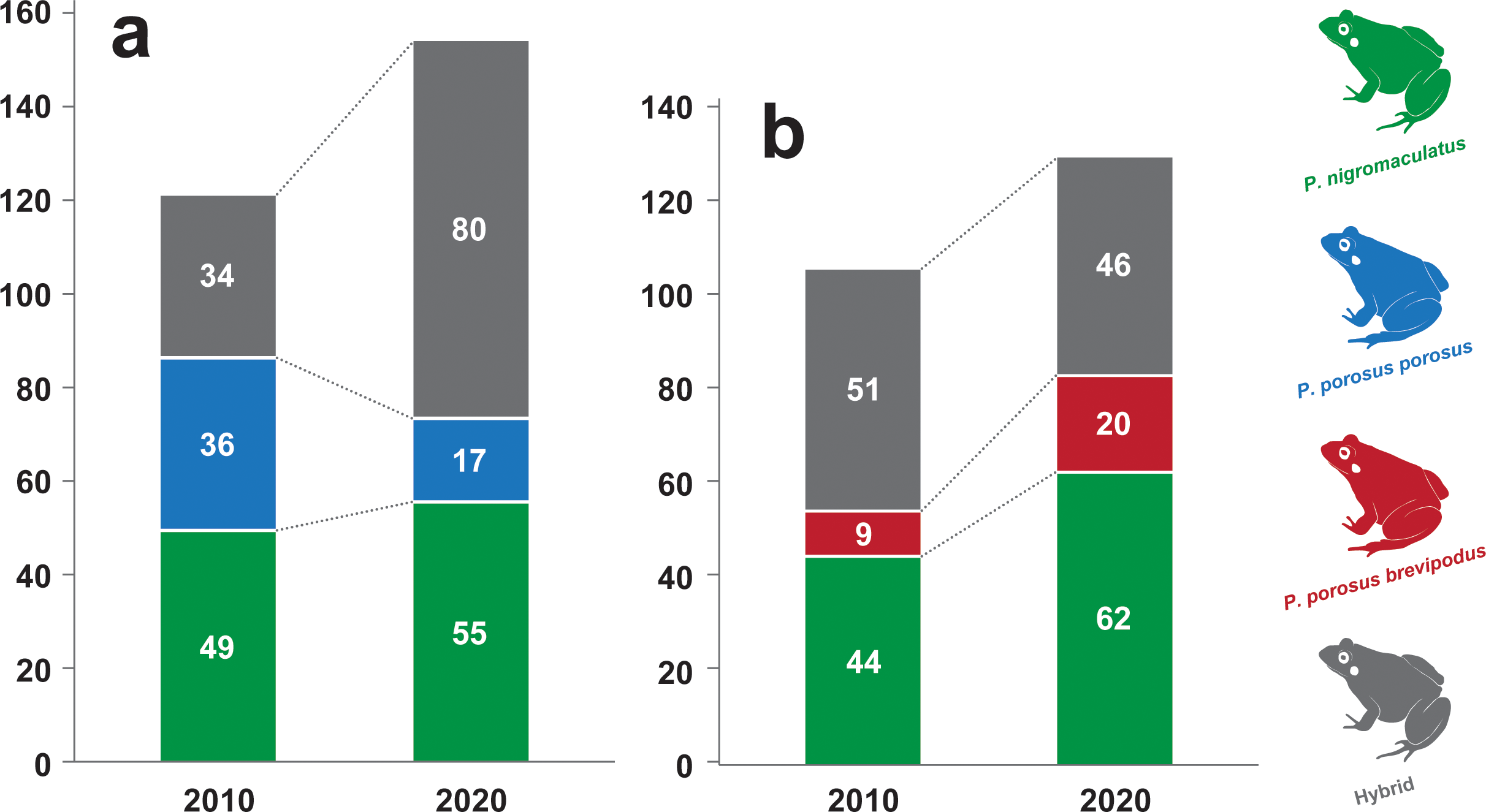
Changes in frequency of pure-bred and hybrid strains detected between 2010 and 2020 of the three analyzed species at survey sites in the Matsumoto Basin (a), and the Ina Basin (b), Nagano Prefecture, Central Japan.

## 4. Discussion

### 4.1 Changes in the distribution pattern of *Pelophylax* frogs in the Matsumoto Basin

Regarding the distribution pattern of *Pelophylax porosus porosus* in the Matsumoto Basin, a slight shrinkage and fragmentation of areas inhabited by pure-bred *P. p. porosus* after the 2010 survey was observed (Komaki et al., 2012). In the survey conducted 10 years ago, many pure-bred *P. p. porosus* were observed in areas where paddy fields were widely and continuously arranged, and there were few roads and building areas that greatly divided the paddy fields. Although many pure-bred *P. p. porosus* were also detected from these areas in the present survey, the proportion of *Pelophylax nigromaculatus* hybrid individuals has increased. In particular at study site 12, since no pure-bred *P. p. porosus* were observed, and replacement with pure-bred *P. nigromaculatus* occurred (Fig. 2, 5), further monitoring is necessary. If hybridization becomes relatively widespread and the fitness of hybrid individuals is lower than that of the parental lineages, wasted reproductive effort may lead to a scenario where the growth rate of one or both populations falls below the rate of hybrid formation, potentially resulting in extinction—a process known as demographic swamping (Wolf et al., 2001; Todesco et al., 2016).

In fact, as mentioned in the “Introduction” section, it is known that F_1_ hybrid males of both species are sterile. In addition, backcrossing with F_1_ females allows hybrid strains to survive, thus backcrossing with the larger body sized *P. nigramaculatus* may give an advantage to hybrid or pure-bred *P. nigramaculatus* strains that are larger and have greater jumping ability (and likely greater dispersal ability) than *P. porosus*. By investigating the frequency of interspecific hybridization and backcrossing, as well as the ecological interspecific interactions that drive these processes, it would be possible to infer the evolutionary fate of the endangered species and provide valuable insights for evolutionary biology and conservation.

Both *P. nigromaculatus* and *P. porosus* have traditionally been thought to be temporally reproductively isolated due to differences in their breeding seasons (Moriya, 1960; Nakanishi et al., 2020). However, their original habitats, the river floodplains, have been rapidly decreasing, and in recent years rice paddy fields have served as alternative habitats. Unfortunately, the amount of rice paddy fields has also decreased largely, and the more recent standardization of agricultural practices has resulted in the period during which rice paddies are flooded being shorter and more limited (i.e., standardization of the flooded periods, mid-season drainage that harms tadpoles due to the introduction of an agricultural practices that involves temporary drainage and drying of rice paddies to eliminate weeds. As such, the temporal separation between *P. nigromaculatus* and *P. porosus* has been eliminated (Serizawa, 1983). As the probability of encounters between the two species, which mainly use paddy fields as breeding grounds, increases, potential reproductive interference between the endangered frogs may increase.

### 4.2 Changes in the distribution pattern of *Pelophylax* frogs in the Ina Basin

In the Ina Basin, interspecific hybridization of *Pelophylax* frogs progressed significantly in the 30 years from the survey 40 years ago to the survey 10 years ago (Shimoyama, 1986, 1999, 2000). Ten years ago, interspecies hybridization was already in serious trouble (Komaki et al., 2012). Therefore, it was feared that further interbreeding of *Pelophylax* frogs would occur over this decade, leading to a decline in pure-bred *P. p. brevipodus*. However, no major changes over the last decade were detected in this study.

In the 30 years from the 1980s to the 2010s, although the area of paddy fields decreased largely, and the number of *Pelophylax* frogs decreased, not much has changed in terms of the genetic characteristics of *Pelophylax* frogs over the last 10 years (Figs. 3, 5). This may be because the paddy field environment has remained relatively unchanged over the last 10 years.

However, the issue of interspecific hybridization is still a conservation concern; it is expected that the proportion of individuals who have experienced interspecific hybridization will increase over time (i.e., change over generations), so the future situation remains unpredictable. Studies of the on genetic structures of *Pelophylax* frogs in both basins and continued conservation efforts for each pure-breed strain are essential.

In both the Matsumoto and Ina Basins, although the genetic hybridization between two *Pelophylax* frog species proceeded rapidly in the 30 years following the survey 40 years ago (Shimoyama, 1986) to the survey 10 years ago (Komaki et al., 2012), the changes in the last 10 years from 2010 to 2020 have not been as drastic as initially feared.

However, local populations consisting of only pure-bred *P. p. porosus* or pure-bred *P. p. brevipodus* do not exist in any of the basins, and it is difficult to avoid further increases in the proportion of hybrid individuals. In addition, pure-bred *P. nigromaculatus* and hybrid strains of *P. nigromaculatus* and *P. p. porosus* were also confirmed at newly added survey sites along the Sai River (No. 29, 30; Fig. 2).

Since these newly added study sites were thought to be inhabited only by pure-bred *P. p. porosus*, it is necessary to examine the expansion of the interspecific hybrid area itself in the future.

Furthermore, the two *Pelophylax* species (two subspecies) targeted in this study are both Red List species. *Pelophylax* frogs’ natural habitat, river floodplains, have been largely lost due to man-made river improvements and other artificial activities, and they have switched to paddy fields as an alternative habitat. However, the number of these paddy fields has been decreasing over the last few decades, and all three *Pelophylax* frog species (including two subspecies) are under threat due to the conversion of waterways to concrete and the standardization of agricultural practices to develop land. In addition to the severity of their habitat situation; moreover, the pure-bred lineages of both species/subspecies are about to be lost due to crossbreeding between the species.

Since this study has visualized areas where pure-bred individuals of each *Pelophylax* frog species have been confirmed, it is also important to conserve the environment in the areas pure-bred individuals inhabit, as well as the pure-bred individuals themselves.

### 4.3 Relationship between changes in frequency of interspecific hybridization and environmental change in both basins

In the Matsumoto and Ina Basins, the rapid expansion of the distribution of *Pelophylax nigromaculatus* and the promotion of interspecific hybridization with *Pelophylax porosus* over the 30 years from the 1980s to 2010 are thought to be due to the decrease in the rice paddy field that function as a substitute for natural floodplains (Komaki et al., 2012). The results of this land use survey also support this trend. In the Matsumoto Basin, even during the 10-year period from 2010 to 2020, when land use changes progressed at a similar pace, there was a trend toward a decrease in the pure-bred *P. p. porosus* individuals and an increase in the proportion of hybrid individuals was observed. However, on the other hand, in the Ina Basin, where land use is also subject to strict conditions, the proportion of purebred *P. p. brevipodus* has remained at relatively similar levels to those observed in 2010. This is considered to be because their habitats in the limited areas where pure-bred *P. p. brevipodus* remain has been maintained in relatively stable conditions for the past 10 years.

Regarding precipitation, no trends were detected over the 50 years surveyed, making it difficult to assess its influence on this Japanese pond frog species. Regarding the temperature data, although a trend over a 50-year period was detected, the respective effects of temperature on the Japanese pond frogs, i.e., *P. nigromaculatus* and *P. porosus*, are not well understood, so evaluating that relationship is a future challenge. In any case, we believe it is important to continue long-term monitoring of the respective population and genetic structureses of the Japanese *Pelophylax* frogs in these two basins over a span of approximately 10-years intervals.

### 4.4 Introgressive frogs with the mtDNA of *P. p. porosus* detected in the Ina Basin

Traditionally, *P. p. porosus* has been considered to be a subspecies endemic to Northeastern Japan, and *P. p. brevipodus* has been considered to be a subspecies endemic to Southwestern Japan (Shimoyama, 1986; Suzuki et al., 2002; Komaki et al., 2012; Matsui and Maeda, 2018). The boundary between these two subspecies corresponds to the western edge of the “Fossa Magna” region (i.e., Itoigawa-Shizuoka Tectonic Line), which is closely related to the formation history of the Japanese Archipelago (Fig. 1). There are many known cases in which closely related species or strains within a species are parapatrically distributed and have this area as their boundary, and which exhibit a close relationship with geological history (Tojo et al., 2017, 2021; Okamiya et al., 2018; Ohba et al., 2020). However, since there is a relatively wide range in the estimated divergence dates of individual cases, it is difficult to understand whether the genetic differentiation observed in such cases was due to the early formation of the Japanese archipelago or the straits in the “Fossa Magna” region, or due to the formation of mountains after the formation of the archipelago. As such, detailed investigations should be conducted in the future in order to clarify the basis of differentiation in such cases. Among the pond frogs that were the focus of this study, *P. nigromaculatus* is certain to have arrived in the Japanese Archipelago later than *P. porosus* (Komaki et al., 2015).

The subspecies differentiation of *P. porosus* is estimated to have occurred around 1.35 Ma, which corresponds to the time when landification of the “Fossa Magna” region was occurring (Komaki et al., 2015). The distribution ranges of both subspecies targeted in this study are separated by a spinal mountainous divide resulting from uplift, which separates the basins located on the Pacific Ocean and Sea of Japan sides. Even today, the topographical division between the Matsumoto Basin and the Ina Basin remains unchanged, and such geohistorical features are considered important factors. Due to these background factors, it is considered that the two subspecies of *P. p. porosus* and *P. p. brevipodus* have maintained their respective populations locally without overlapping each other’s distribution areas.

In this study, mtDNA sequences of *P. p. porosus* were detected from three individuals (2.3%) out of 128 specimens collected in the Ina Basin and genetically analyzed. Conversely, no mtDNA sequences of *P. p. brevipodus* were detected from 158 specimens collected in the Matsumoto Basin and genetically analyzed (Figs 2, 5).

Although it is possible that there was an introgression between the two subspecies of *P. porosus*, such introgression did not occur near the boundary between the distribution ranges of the two subspecies. *Pelophylax porosus* gene flow of both subspecies did not occur near the boundaries of their distribution ranges, and these three individuals were all confirmed from point 28 in the most southerly part of the Ina Basin (Figs 3, 5). If the distributions of the two subspecies have overlapped in the past, it would not be surprising to see signs of inter-subspecies hybridization more frequently.

Another possibility is that interspecific hybrids (or their progeny) of *P. p. porosus* (female) and *P. nigromaculatus* (male) within the Matsumoto Basin moved and dispersed to the Ina Basin. The hypothesis that females of such hybrid lines enter the Ina Basin while repeatedly backcrossing with *P. nigromaculatus* males may explain why only a few individuals in the Ina Basin have *P. p. porosus* mtDNA. In fact, *P. nigromaculatus* has a larger body size than *P. porosus*, and it is thought to have a higher jumping ability and dispersal ability (Maeda and Matsui, 1999), so we think this is clearly possible.

If these hypotheses are correct, *P. p. porosus* and *P. p. brevipodus* never had direct contact with each other, and a third party, *P. nigromaculatus*, functioned as a “genome vehicle”. Therefore, this is a very interesting phenomenon because it is as if the genome itself has been “hitchhiking”. However, the possibility of artificial dispersion or the possibility of temporary large-scale flooding events that resulted in accidental long-distance dispersal to downstream river areas cannot be discounted. In any case, expanding the survey area and conducting studies focusing on recent gene flow may reveal the presence of *P. p. porosus* type DNA within Ina Basin. In the future, we would like to confirm hypothesis this by conducting genome-wide analyses using more sensitive gene markers, e.g., SSR, MIG-seq and GRAS-Di analyses.

## 5. Conclusion

In the central part of the Japanese Archipelago, the distribution ranges of *Pelophylax nigromaculatus* and *Pelophylax porosus* overlap. Our previous study conducted in 2010 revealed that the interspecific hybridization had occurred between *P. nigromaculatus* and *P. p. porosus* in the Matsumoto Basin, and also interspecific hybridization between *P. nigromaculatus* and *P. p. brevipodus* in the Ina Basin. A similar survey was conducted in these regions in 2020, 10 years after the earlier study, to assess the extent of interspecific hybridization progress over the 10-year period. In evaluating pure lineage individuals and hybrid individuals, we used sequences from the mitochondrial DNA Cyt-b (586-bp) region and nuclear DNA Tyrosinase (747-bp) region. In the Matsumoto Basin in 2010 and 2020, 119 and 171 specimens were genetically analyzed, respectively. In the Ina Basin in 2010 and 2020, 104 and 128 specimens were genetically analyzed, respectively. As a result of this, in the Ina Basin, where there were only a few “pure”-bred *P. p. brevicodus* in 2010, it showed almost the same trend in 2020. Although there are still concerns about the conservation of the *P. p. brevipodus* lineage, no progress in hybridization has been observed over the past 10-year period. On the other hand, in the Matsumoto Basin, interspecific hybridization between *P. nigromaculatus* and *P. p. porosus* was evaluated to have progressed slightly. Both of these species are listed on the Red List by the Ministry of the Environment and also by local governments (i.e., Nagano Prefecture), and there are concerns that the number of purebred individuals of each species will decrease, so continued monitoring will be necessary.

## Acknowledgments

We are grateful to Doctors Atsushi Iwasawa (Gifu University), Koki Yano, Seiya Okamoto, Gaku Ueki (Shinshu University), and Masters students Takahisa Ozaki, Kanako Momose, Haruka Yamazaki, Riko Kawauchi, Keisuke Inoue, Ryota Tomizawa (Shinshu University) for their valuable comments. This study was supported by grants from the Chikuma-gawa Research Group of the River Ecology Research Organization and the Institute of Mountain Science, Shinshu University. This study was conducted in accordance with the ethical regulations of Shinshu University.

## Funding

This study was supported by a research grant from the Institute of Mountain Science, Shinshu University (KT).

## Contributions

S. S., T. S. and K. T. designed the research; S. S. performed the field experiments; S. S. and T. S. perdormed the DNA analyzed and sequenced data analyzed; and all authors wrote the original draft of the paper.

## Data Availability Statement

The nucleotide sequence data generated in this study are uploaded as supplementary material for review only and are not currently deposited in GenBank. The data will be deposited in GenBank upon publication, and accession numbers will be provided in the final version. The location information associated with these sequences is explicitly provided in Table 1 of the manuscript.

## Table Captions

**Table S1** The results of calculating the area by land use type in each basin in three years are shown. The data used was from the GIS data provided by the Ministry of Land, Infrastructure, Transport and Tourism (https://nlftp.mlit.go.jp/ksj/index.html) for the year closest to the survey year. (a) Shows the area of the three most occupied land use types, excluding forest, for each basin and year. (b) Shows the calculated area of all land use types for each basin and year.

## Figure Legends

**Figure S1** Environmental analyses for the two basins over the last few decades, i.e., temperature and precipitation. Fifty years of observational data of temporal variations in temperature and precipitation from 1970 to 2020 in the Matsumoto (Matsumoto Meteorological Observatory) and Ina Basins (Iida Meteorological Observatory). These analyses of temporal variations using the STL decomposition method. Analyses of temporal variations in temperature and precipitation in the Matsumoto Basin and Ina Basin from 1970 to 2020 using the STL decomposition method. The data, trend components, seasonal components, and residual components are shown for (a) the daily maximum temperature in the Matsumoto Basin, (b) the daily maximum temperature in the Ina Basin, (c) the daily minimum temperature in the Matsumoto Basin, (d) the daily minimum temperature in the Ina Basin, (e) the daily mean temperature in the Matsumoto Basin, (f) the daily mean temperature in the Ina Basin, (g) the monthly precipitation in the Matsumoto Basin, and (h) the monthly precipitation in the Ina Basin. The top row of each figure shows the original data and trend components, the middle row shows the seasonal components, and the bottom row shows the reminder components. The box highlights in the figures indicate the survey years, with 1980, 2010, and 2020 highlighted, respectively.

It became clear that due to the effects of global warming and/or urbanization (described below), the trends in the monthly average daily maximum temperatures, monthly average daily minimum temperatures, and monthly average daily temperatures in both basins have all been gradually rising over the past 50 years. Precipitation varies greatly within each year, with no clear trends observed over the past 50 years.

**Figure S2** Environmental analyses for the two basins over the last few decades, i.e., land use changes from 1987 to 2021 in the both basins. (a) the Matsumoto Basin in 1987, (b) the Matsumoto Basin in 2009, (c) the Matsumoto Basin in 2021, (d) the Ina Basin in 1987, (e) the Ina Basin in 2009, (f) the Ina Basin in 2021.

The results are shown color-coded by land use type for each basin and each year. Land use types are assigned according to the priority of features in each 100 m mesh. Regarding land use, it was revealed that the area of paddy fields had decreased significantly over the 22 years from 1987 to 2009 (a 15.1% decrease in the Matsumoto Basin and a 17.4% decrease in the Ina Basin). In addition, although the decrease over the 12 years from 2009 to 2021 was less (a decrease of 5.2% in the Matsumoto Basin and 8.5% in the Ina Basin), which when averaged gives an annual rate of decrease that has been almost constant. Meanwhile, as the area of farmland and building areas has increased, it is believed that over the past 35 years there has been a great deal of land conversion from paddy fields to farmland and residential land, and from paddy fields to farmland and then residential land.

## Notes

### Competing Interest Statement

The authors have declared no competing interest.

## References

Bossuyt, F., and M.C. Milinkovitch. 2000. Convergent adaptive radiations in Madagascan and Asian ranid frogs reveal covariation between larval and adult traits. Proceedings of the national Academy of Sciences 97(12): 6585−6590. 10.1073/pnas.97.12.6585

Cleveland, R. B., W. S. Cleveland, J. E. McRae, and I. Terpenning. 1990. STL: A seasonal-trend decomposition. Journal of Official Statistics 6(1): 3–73. https://www.nniiem.ru/file/news/2016/stl-statistical-model.pdf

Kameyama, T., T. Morita, S. Okada, J. Naito, and T. Utsunomiya. 2006. Crisis and reintroduction of the Okayama race of the Daruma pond frog, Rana porosa brevipoda. Japanese Journal of Conservation Ecology 11: 158−166. 10.18960/hozen.11.2_158

Katano, O., K. Hosoya, K. I. Iguchi, M. Yamaguchi, Y. Aonuma, and S. Kitano. 2003. Species diversity and abundance of freshwater fishes in irrigation ditches around rice fields. Environmental Biology of Fishes 66: 107–121. 10.1023/A:1023678401886

Komaki, S., T. Igawa, S. M. Lin, K. Tojo, M. S. Min, and M. Sumida. 2015. Robust molecular phylogeny and palaeodistribution modelling resolve a complex evolutionary history: glacial cycling drove recurrent mt DNA introgression among *Pelophylax* frogs in East Asia. Journal of Biogeography, 42(11): 2159–2171. 10.1111/jbi.12584

Komaki, S., A. Kurabayashi, M. M. Islam, K. Tojo, and M. Sumida. 2012. Distributional change and epidemic introgression in overlapping areas of Japanese pond frog species over 30 years. Zoological Science 29(6): 351–358. 10.2108/zsj.29.351

Komaki, S., and K. Tojo. 2010. Distribution patterns of the closely related three Japanese pond frogs in the Matsumoto and Ina Basins (Nagano Prefecture), and the possibility of interspecific hybridization in the overlapping areas of their distributions. The Bulletin of Shiojiri City Museum of Natural History 11: 52–60.

Kumar, S., G. Stecher, and K. Tamura. 2016. MEGA7: molecular evolutionary genetics analysis version 7.0 for bigger datasets. Molecular Biology and Evolution 33(7): 1870–1874. 10.1093/molbev/msw054

Leigh, J. W., D. Bryant, and S. Nakagawa. 2015. POPART: full-feature software for haplotype network construction. Methods in Ecology & Evolution, 6(9): 1110–1116. 10.1111/2041-210X.12410

Maeda, N., and M. Matsui. 1999. Frogs and Toads of Japan, Revised edition. Bun-ichi Sogo Shuppan, Tokyo

Matsui, M., and N. Maeda. 2018. Encyclopaedia of Japanese frogs. Bun-ichi Sogo Shuppan, Tokyo

Ministry of the Environment, Government of Japan. 2020. *Red* List of Japan. (in Japanese.) Available from URL: https://ikilog.biodic.go.jp/Rdb/booklist

Moriya, K. 1960. Studies on the five races of the Japanese pond frogs, *Rana nigromaculata* Hallowell III. Sterility in interracial hybrids. Journal of Science Hiroshima University Series B, Division 1. Zoology 18: 109–124.

Naito, R. 2012. Perspectives of conservation of pond-breeding frogs (focusing on the Nagoya Daruma pond frog) in rice paddy areas in Japan. Landscape Ecology and Management 17(2): 57–73. 10.5738/jale.17.57

Nakanishi, K., A. Honma, M. Furukawa, K. I. Takakura, N. Fujii, K. Morii, Y. Terasawa, and T. Nishida. 2020. Habitat partitioning of two closely related pond frogs, *Pelophylax nigromaculatus* and *Pelophylax porosus brevipodus*, during their breeding season. Evolutionary Ecology, 34, 855–866. 10.1007/s10682-020-10061-1

Nishioka, M., M. Sumida, and H. Ohtani. 1992. Differentiation of 70 populations in the *Rana nigromaculata* group by the method of electrophoretic analyses. Scientific Report of the Laboratory for Amphibian Biology, 11, 1–70.

Ohba, S. Y., K. Numata, and K. Kawano (2020). Variation in flash speed of Japanese firefly, *Luciola cruciata* (Coleoptera: Lampyridae), identifies distinct southern “quickLflash” population on Goto Islands, Japan. Entomological Science, 23(2), 119–127. 10.1111/ens.12403

Okamiya, H., H. Sugawara, M. Nagano, and N. A. Poyarkov. 2018. An integrative taxonomic analysis reveals a new species of lotic *Hynobius* salamander from Japan. PeerJ, 6: e5084. 10.7717/peerj.5084

Rozas, J., A. Ferrer-Mata, J. C. Sánchez-DelBarrio, S. Guirao-Rico, P. Librado, S. E. Ramos-Onsins, and Sánchez-Gracia, A. (2017). DnaSP 6: DNA sequence polymorphism analysis of large data sets. Molecular Biology and Evolution, 34(12): 3299–3302. 10.1093/molbev/msx248

Sagiya, T., T., Nishimura, Y. Iio, and T. Tada. 2002. Crustal deformation around the northern and central Itoigawa-Shizuoka Tectonic Line. Earth, Planets and Space, 54(11): 1059–1063. 10.1186/BF03353302

Serizawa, T. 1983. Reproductive traits of the *Rana nigromaculata*-*brevipoda* complex in Japan. I. Growth and egg-laying in Tatsuda and Saya, Aichi prefecture. Japanese Journal of Herpetology, 10: 7–19.

Shimoyama, R. 1986. Distribution and life history of the *Rana nigromaculata* species group in Nagano Prefecture, Japan. Shinano Kyoiku 1199: 68–73 (in Japanese)

Shimoyama, R. 1999. Interspecific interactions between two Japanese pond frogs, *Rana porosa brevipoda* and *Rana nigromaculata*. Japanese Journal of Herpetology, 18(1): 7–15. 10.5358/hsj1972.18.1_7

Shimoyama, R. 2000 Conspecific and heterospecific pair-formation in *Rana porosa brevipoda* and *Rana nigromaculata*, with reference to asymmetric hybridization. Current herpetology19(1): 15–26. 10.5358/hsj.19.15

Suzuki, K., K. Okubo, and T. Sawahata. 2002. The distribution and population density of the threatened species *Rana porosa brevipoda* and conditions of paddies as frog habitats in the Ina basin, Nagano prefecture, in central Japan. Journal of the Japanese Institute of Landscape Architecture (Japan), 65(5): 517–522. 10.5632/jila.65.517

Suzuki, T., K. Yano, S. Okamoto, G. Ueki, A. Fukakusa, M. Ikeda, G. Inoue, H. Tagashira, T. Yoshida, and K. Tojo. 2023. A major flood caused by a typhoon did not affect the population genetic structure of a river mayfly metapopulation. Proceedings of the Royal Society B, 290 (1997), 20230177. 10.1098/rspb.2023.0177

Tanaka, T., M. Matsui, and O. Takenaka. 1996. Phylogenetic relationships of Japanese brown frogs (*Rana*: Ranidae) assessed by mitochondrial cytochrome b gene sequences. Biochemical Systematics and Ecology 24(4): 299–307. 10.1016/0305-1978(96)00017-8

Todesco, M., M. A. Pascual, G. L. Owens, B. T. Ostevik, S. Moyers, …, and L. H. Rieseberg, L. H. (2016). Hybridization and extinction. Evolutionary applications, 9(7), 892–908. 10.1111/eva.12367

Tomita, K., T. Suzuki, K. Yano, and K. Tojo. 2020. Community structure of aquatic insects adapted to lentic water environments, and fine-scale analyses of local population structures and the genetic structures of an endangered giant water bug *Appasus japonicus*. Insects 11(6): 389. 10.3390/insects11060389

Tojo, K., K. Sekiné, M. Takenaka, Y. Isaka, S. Komaki, S. Suzuki, and S. D. Schoville. 2017. Species diversity of insects in Japan: their origins and diversification processes. Entomological Science 20(1): 357–381. 10.1111/ens.12261.

Tojo, K., K. Miyairi, Y. Kato, A. Sakano, and T. Suzuki. 2021. A description of the second species of the genus *Bleptus* Eaton, 1885 (Ephemeroptera: Heptageniidae) from Japan, and phylogenetic relationships of two *Bleptus* mayflies inferred from mitochondrial and nuclear gene sequences. Zootaxa 4974(2): 333360–333360. 10.11646/zootaxa.4974.2.5

Wolf, D. E., N. Takebayashi, and L. H. Rieseberg. 2001. Predicting the risk of extinction through hybridization. Conservation Biology, 15(4), 1039–1053. 10.1046/j.1523-1739.2001.0150041039.x

Yamamoto, Y., and Y. Senga (2012). The distribution of the Tokyo Daruma pond frog, *Rana porosa porosa*, and its habitat status in paddy fields fragmented by urbanization. Japanese Journal of Conservation Ecology 17: 175–184. 10.18960/hozen.17.2_175

Yamazaki, M., Y. Hamada, T. Ibuka, T. Momii, and M. Kimura 2001. Seasonal variations in the community structure of aquatic organisms in a paddy field under a long-term fertilizer trial. Soil Science and Plant Nutrition 47(3): 587–599. 10.1080/00380768.2001.10408422

