## Supplementary figures and images for "Changes over a 10-year Period in the Distribution Ranges and Genetic Hybridization of Three *Pelophylax* Pond Frogs in Central Japan"

### Figure S1

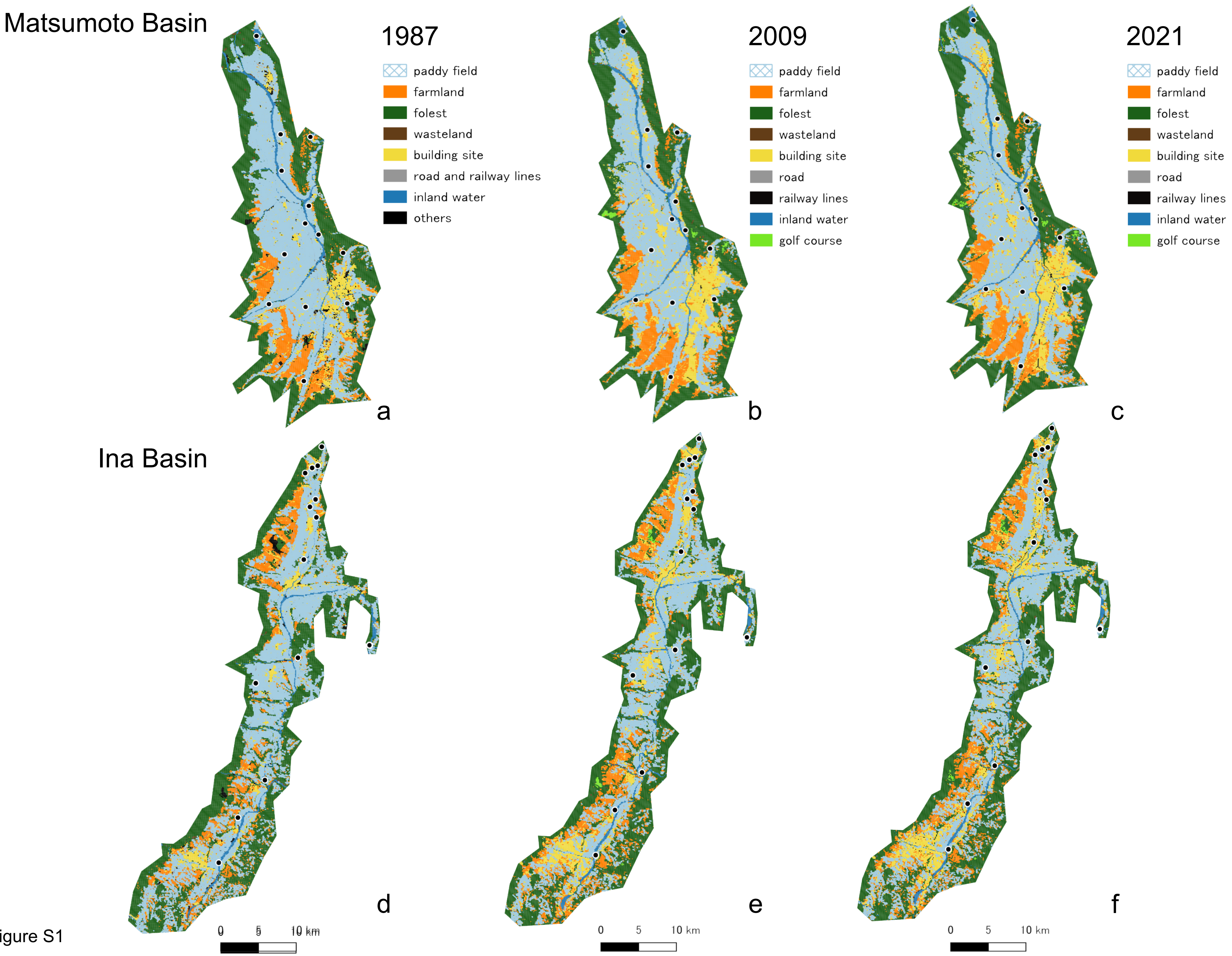

### Figure S2

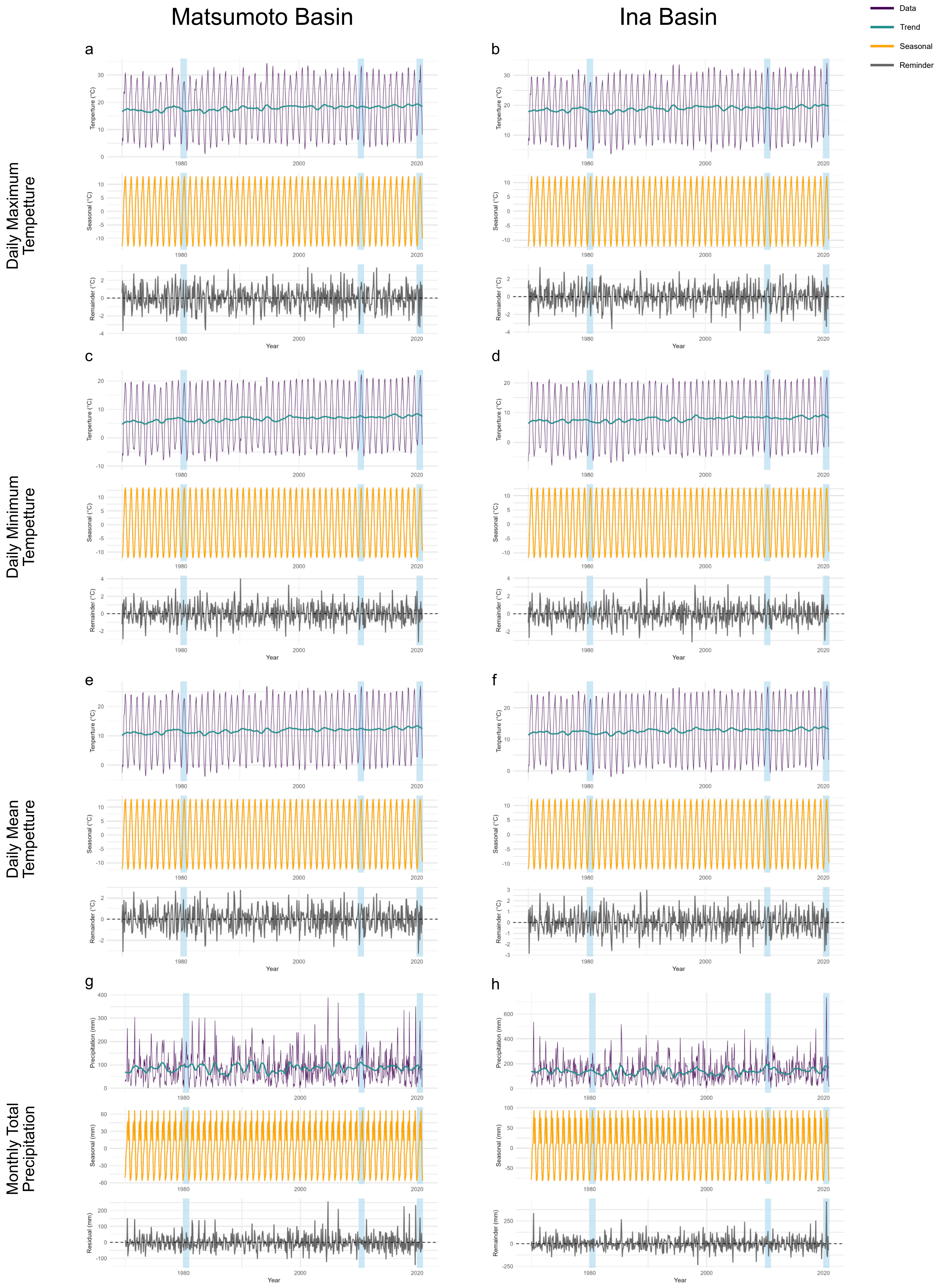
