## Supplementary material for "Changes over a 10-year Period in the Distribution Ranges and Genetic Hybridization of Three *Pelophylax* Pond Frogs in Central Japan": Table S1

Table S1 The results of calculating the areas by land use type of the both basin in each year of 1987, 2009 and 2021 are shown. The data used was from the GIS data provided by the Ministry of Land, Infrastructure, Transport and Tourism (https://nlftp.mlit.go.jp/ksj/index.html) for the year closest to the survey year. The table shows the land use types with particularly large occupied areas in each basin by year

| | **Matsumoto basin (area, m^2)** | | | **Ina basin (area, m^2)** | | |
|---|---|---|---|---|---|---|
| | **1987** | **2009** | **2021** | **1987** | **2009** | **2021** |
| **Forest** | 224,616,229 | 221,848,325 | 217,942,472 | 257,787,120 | 249,382,965 | 250,389,265 |
| **Paddy field area** | 220,709,742 | 187,314,787 | 177,494,898 | 184,803,633 | 152,678,061 | 139,665,461 |
| **Farmland area** | 91,398,088 | 92,751,464 | 96,907,577 | 102,567,462 | 107,424,849 | 110,806,605 |
| **Building area** | 81,852,533 | 134,585,896 | 137,367,610 | 59,120,539 | 105,409,847 | 109,082,307 |
| **Inlamdwater** | 29,845,690 | 25,944,476 | 29,960,390 | 22,311,566 | 20,424,919 | 22,580,575 |
